# Phylogenomics reveals dynamic evolution of fungal nitric oxide reductases and their relationship to secondary metabolism

**DOI:** 10.1101/301895

**Authors:** Steven A. Higgins, Christopher W. Schadt, Patrick B. Matheny, Frank E. Löffler

## Abstract

Fungi expressing P450nor, an unconventional nitric oxide (NO) reducing cytochrome P450, are thought to be significant contributors to soil nitrous oxide (N_2_O) emissions. However, fungal contributions to N_2_O emissions remain uncertain due to inconsistencies in measurements of N_2_O formation by fungi. Much of the N_2_O emitted from antibiotic-amended soil microcosms is attributed to fungal activity, yet fungal isolates examined in pure culture are poor N_2_O producers. To assist in reconciling these conflicting observations and produce a benchmark genomic analysis of fungal denitrifiers, genes underlying fungal denitrification were examined in >700 fungal genomes. Of 167 *p450nor*–containing genomes identified, 0, 30, and 48 also harbored the denitrification genes *narG*, *napA* or *nirK*, respectively. Compared to *napA* and *nirK*, *p450nor* was twice as abundant and exhibited two to five-fold more gene duplications, losses, and transfers, indicating a disconnect between *p450nor* presence and denitrification potential. Furthermore, co-occurrence of *p450nor* with genes encoding NO-detoxifying flavohemoglobins (Spearman r = 0.87, *p* = 1.6e^−10^) confounds hypotheses regarding P450nor’s primary role in NO detoxification. Instead, ancestral state reconstruction united P450nor with actinobacterial cytochrome P450s (CYP105) involved in secondary metabolism (SM) and 19 (11 %) *p450nor*-containing genomic regions were predicted to be SM clusters. Another 40 (24 %) genomes harbored genes nearby *p450nor* predicted to encode hallmark SM functions, providing additional contextual evidence linking *p450nor* to SM. These findings underscore the potential physiological implications of widespread *p450nor* gene transfer, support the novel affiliation of *p450nor* with fungal SM, and challenge the hypothesis of *p450nor*’s primary role in denitrification.

## Importance

Fungi are considered substantial contributors to emissions of the greenhouse gas N_2_O, owing to the nitric oxide reducing potential of an unusual cytochrome P450 (P450nor). Despite these findings, fungi do not satisfy criteria to be classified as respiratory denitrifiers and methodological biases confound fungal contributions to the N_2_O budget. Phylogenetic and genomic analyses distanced N_2_O-forming fungi from denitrification and supported a new link between P450nor and SM. Hence, N_2_O formed by P450nor activity may be artificially induced or a byproduct of SM. Explorations of P450nor’s involvement in SM may facilitate the discovery of new compounds with potential applications in agricultural and pharmaceutical industries. Dissociating *p450nor* from denitrification also informs climate change models and directs research towards organisms and processes most relevant to *in situ* N_2_O production.

## Introduction

Increased human reliance on fixed nitrogen (N) from the Haber-Bosch process to meet the demands of sustaining an expanding global population has contributed to a 20 % increase in atmospheric nitrous oxide (N_2_O), a potent greenhouse gas with ozone destruction potential (1, 2). N_2_O is primarily formed by denitrifying members of the *Bacteria* and *Archaea* (3), a prevailing view that has been challenged by experiments reporting that abundant soil- and sediment-inhabiting fungi contribute up to 89 % of the total N_2_O emitted from these systems (4–6). Notably, fungi cannot convert N_2_O to inert N2 like many denitrifying bacteria (7), suggesting their contributions to greenhouse effects and ozone destruction could be significant. Fungi are considered to be important sources of N_2_O emissions from agroecosystems (8, 9), which are predicted to contribute up to two-thirds of the total N_2_O emissions by 2030 (10). Studies of model fungi show that N_2_O formation is due to P450nor, a heme-containing cytochrome P450, that catalyzes the two electron reduction of nitric oxide (NO) to N_2_O (11–13). N_2_O formation by P450nor is thought to occur exclusively in fungi and the *p450nor* gene has been exploited as a distinctive biomarker in molecular assays to study fungal denitrifier diversity and abundance in the environment (14–16).

Despite these observations, the fungal contributions to N_2_O emissions remain uncertain. For example, fungi do not satisfy criteria set forth to classify microorganisms as respiratory denitrifiers (17). N_2_O-producing fungi in pure culture do not exhibit a balance between the inorganic N inputs and quantities of N_2_O formed (18–20) and possess three to six orders of magnitude lower rates of N_2_O production compared to denitrifying bacterial isolates under optimal conditions (4). Fungi also fail to generate anoxic growth yields proportional to the quantity of inorganic N reduced in pure culture (6, 21–23), and no significant relationship was detected between fungal denitrification activity and fungal biomass in anoxic soil incubations (24). Above all, partitioning techniques (antibiotic inhibition and isotope site preference) used to estimate fungal and bacterial contributions to N_2_O emissions are biased and often lack corroborating evidence in conjunction with their application, suggesting fungal contributions to N_2_O emissions are substantially inflated (5, 25–27). For example, antibiotics are often criticized for lacking both generality and specificity, but the expected biases resulting from the exclusive use of antibiotic inhibition techniques to assess fungal contributions to N_2_O emissions remain unaccounted for. Bias could be interpreted by concurrently employing culture-independent techniques (i.e., multi-omics approaches); however, these practices are lacking in investigations of fungal denitrification, and the singular use of antibiotics to partition microbial activity casts doubt on the quantitative value of observations derived from this approach regarding fungal denitrification (25, 26).

The capacity for N_2_O-production conferred by *p450nor* in fungi is a uniquely eukaryotic trait, yet previous investigations have hypothesized an actinobacterial origin for *p450nor* based on sequence comparisons (7, 28–30). Of note, *Actinobacteria* are not considered canonical denitrifying bacteria, and only a few reports of their denitrification capacity exist (31–33). Most members of the *Actinobacteria* possess a truncated denitrification pathway or lack a canonical nitric oxide reductase gene (*nor*) (with the exception of *Corynebacterium* and *Propionibacterium*) (32, 33). Hence, members of the Fungi and *Actinobacteria* share an incomplete denitrification pathway with a potentially limited capacity to perform denitrification. Consistent with the horizontal gene transfer (HGT) hypothesis are sequence similarities between fungal P450nor and actinobacterial P450s of the CYP105 family, many of which have been investigated for their contributions to secondary metabolism (SM) (7, 34). Despite these observations, the prevailing hypothesis regarding *p450nor*’s evolution and function was its acquisition from the *Actinobacteria* and subsequent evolution to fill a novel role in denitrification, specifically the reduction of NO to N_2_O (7, 29). The hypothesis that *p450nor* was acquired from one or more members of the *Actinobacteria* and retained an ancestral function in SM surprisingly remains unexplored.

Efforts associated with the 1,000 Fungal Genomes and Assembling the Fungal Tree of Life (AFTOL) projects have resulted in a steady rise in genomic sequence data for members of the fungal kingdom (35, 36). These large scale sequencing efforts facilitate comprehensive phylogenomic investigations with the potential to uncover the causes and consequences of the genomic architecture of fungi and assist in directing research efforts. Hence, the overarching questions this study addresses are I) what is the breadth of denitrification genes across fungal genomes and what are their evolutionary relationships, and II) can phylogenomic analyses reconcile the conflict in fungal contributions to N_2_O formation observed in laboratory and environmental settings? Our comparative genomic and phylogenetic analyses identified a disconnect between *p450nor* and denitrification gene presence and supported a role for P450nor in SM rather than denitrification. Importantly, these results provide an explanation for the minor, non-respiratory capacity of fungi to form N_2_O, and suggests N_2_O is a byproduct of active SM. These findings transform our understanding of the ecological significance and environmental consequences of *p450nor* presence/absence in fungal genomes.

## Results

### Infrequent co-occurrence among denitrification genes in fungi

Bioinformatic analyses identified homologs of canonical bacterial and fungal denitrification genes (*narG*, *napA*, *norB*, *nirK*, *nosZ*, *p450nor*) in 712 fungal genomes. Of the denitrification gene set investigated, only *narG*, *napA*, *nirK*, and *p450nor* were detected (Fig. 1). Genes encoding the membrane bound respiratory nitrate reductase (*narG*) were detected in only three fungal genomes (0.42 %) and were excluded from further analysis due to their low occurrence. The genes predicted to encode the periplasmic nitrate reductase (NapA) and the copper-containing nitrite reductase (NirK) were detected in 75 (10.5 %) and 82 (11.5 %) of the 712 fungal genomes analyzed, respectively (Fig. 1, Table S1). In contrast, P450nor gene sequences occurred at approximately twice the frequency in 167 (23 %) of the fungal genomes analyzed, supporting the claim that P450nor-mediated N_2_O production may be widespread in fungi (Fig. 1) (37). A breakdown of genus- and family-level denitrification gene abundances in fungal genomes underscores the disparity in presence/absence of denitrification genes in fungi and is available in Supplemental Information (SI) (Dataset S1, Fig. S1).

**Figure 1.**
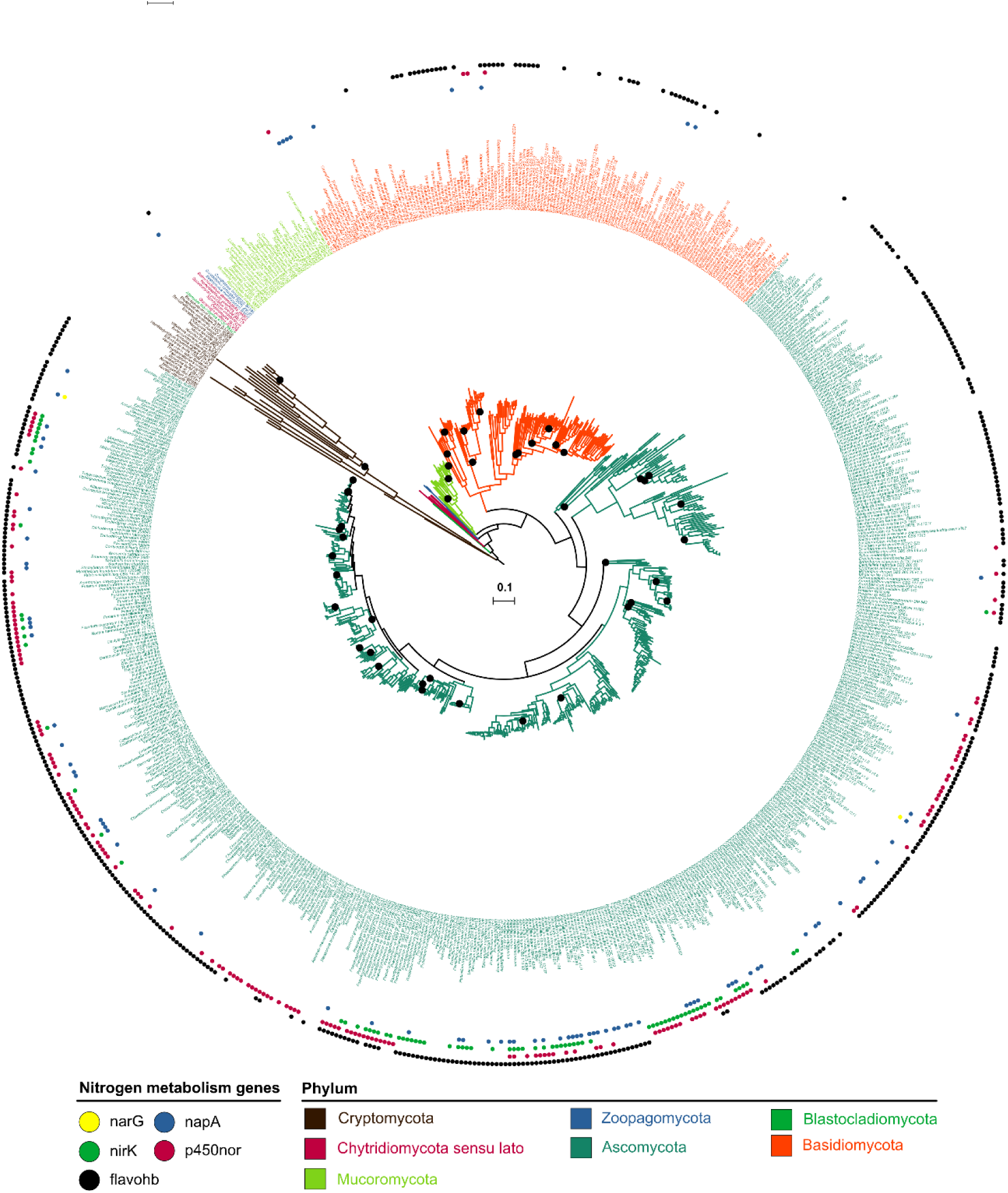
Maximum-Likelihood phylogeny of the kingdom Fungi inferred from a concatenated alignment of 238 single copy marker gene amino acid sequences (see Materials and Methods). Black circles marking branches indicate nodes with bootstrap percentages below 90%. Colored markers outside taxon names specify the presence or absence of each gene (narG, napA, nirK, p450nor, flavohemoglobin) within a fungal genome. Flavohb, flavohemiglobin genes involved in NO detoxification. The scale bar (center of tree) represents amino acid substitutions per site. A high-resolution file of the tree is available at https://doi.org/10.6084/m9.figshare.c.3845692.

Our analyses also revealed a low co-occurrence between *p450nor* and additional fungal denitrification pathway markers. Since *p450nor* is regarded as the sole trait encoding N_2_O production in fungi, the co-occurrence of multiple denitrification gene markers would be indicative of a capacity for sequential respiratory denitrification, whereas isolated occurrences could be indicative of alternative processes such as detoxification. The three-gene set *narG*/*nirK*/*p450nor* did not co-occur in any of the fungal genomes examined, whereas co-occurrence of the gene set *napA*/*nirK*/*p450nor* was observed in 18 (10.8 %) of 167 *p450nor*-containing fungal genomes. Sets of at least two co-occurring denitrification traits (i.e., *narG*/*p450nor*, *napA*/*p450nor*, and *nirK*/*p450nor*) were found in 0, 18 and 29 % of fungal genomes, respectively. Of the *napA*-containing fungal genomes, 25 (33 %) also contained a *nirK* gene, whereas 30 % of the *nirK*-containing fungal genomes also harbored a *napA* gene. Evolutionary correlation was strongly supported for the gene sets *napA*/*nirK*, *napA*/*p450nor*, and *nirK*/*p450nor*, with average log Bayes Factor values of 31.9 ± 0.60, 12.2 ± 0.11, and 31.3 ± 0.04, respectively. Hence, the genes *napA*, *nirK*, and *p450nor* occur in related fungal taxa, but co-occurrences were infrequent within the individual fungal genomes analyzed.

### Evolutionary forces acting upon denitrification traits within fungi

To identify evolutionary forces shaping the observed distribution of denitrification traits within fungi, comparisons between gene and species trees were assessed with phylogenetic tests and parsimony-informed models to quantify evolutionary events. Visual inspection of *p450nor* gene and species trees indicated potential widespread HGT of *p450nor* within fungi, examples of which included HGT of *p450nor* from the phylum Ascomycota to members of the Basidiomycota and within and among classes of ascomycetes (Fig. S2). Furthermore, the monophyly of five fungal classes containing *p450nor* (Dothideomycetes, Eurotiomycetes, Leotiomycetes, Sordariomycetes, and Tremellomycetes) were not supported by approximately unbiased (AU) tests (*p* ≤ 0.05, Table S2), indicative of dynamic evolution of *p450nor* in most fungal lineages. Although co-phylogeny plots are suggestive of HGT, additional analysis using NOTUNG software was performed to model potential gene duplication (GD), gene transfer (GT), and gene loss (GL) events (38). Of the *napA*, *nirK*, and *p450nor* genes analyzed, the *p450nor* phylogenies had the greatest number of predicted GT events, ranging from 4 to 15 GT events despite applying stringent GT costs within NOTUNG software (Table S3). At GT costs below 9, no temporally consistent optimal solutions were reached, suggesting that GD and GL alone are insufficient to describe the evolutionary dynamics of *p450nor* in fungi. Using the same stringent GT costs, the predicted number of GT events detected for *napA* and *nirK* were much lower, and ranged from 1 to 3 and 0 to 1 GT events for each gene, respectively (Table S3). The reduced number of GT events detected in *napA* and *nirK* phylogenies were also apparent from co-phylogeny plots of each gene (Fig. S3, S4) compared to co-phylogenetic plots for *p450nor* (Fig. S2). Although GT events detected for *napA* were lower than *p450nor* at high GT costs, GT may still represent a significant evolutionary force contributing to the observed *napA* distribution in extant fungal lineages (Table S3). For example, AU tests rejected the monophyly of three Ascomycota (Dothideomycetes, Leotiomycetes, and Sordariomycetes) and one Basidiomycota (Pucciniomycetes) lineage within the *napA* phylogeny (Table S2, P ≤ 0.05). Specific instances of predicted HGT events are outlined in Supplemental Information (SI) for each gene (Table S4, Fig. S5).

### Fungal P450nor evolved from actinobacterial P450s involved in SM

Previous investigations have hypothesized an actinobacterial origin for *p450nor* based on amino acid sequence alignments (7, 28, 30), but rigorous phylogenetic tests of *p450nor*’s origins were lacking to support this hypothesis. Alignment of fungal P450nor amino acid sequences to the NCBI RefSeq protein database identified 230 bacterial sequences with significant sequence alignment (≥ 65 % query coverage, ≥ 35 % amino acid identity) to P450nor. Of note, *p450nor* homologs were also detected within the genomes of three freshwater inhabiting green algae, *Chlorella variabilis*, *Chlamydomonas reinhardtii*, and *Monoraphidium neglectum*, expanding the known distribution of *p450nor* to photosynthetic eukaryotic microbes. Additional *p450nor* homologs were not detected in archaea, plant, protist, or other lineages housed within the RefSeq database. Of the bacterial cytochrome P450 (hereafter P450) sequences identified, approximately 6 % (n = 13) were proteobacterial in origin, whereas the remaining sequences belonged to members of the bacterial phylum *Actinobacteria* (Fig. S6). Ancestral character state reconstruction of select P450 families on a subset of these sequences supported the monophyly of *p450nor* and bacterial P450 gene sequences of the P450 family CYP105 (Fig. 2) (39). The same relationships were preserved when phylogenetic reconstruction was performed using the complete set of 408 P450 amino acid sequences (Fig. S7). Importantly, NO-utilizing P450 sequences from the CYP107 family belonging to members of the *Streptomyces* formed a larger monophyletic clade containing P450nor and other CYP105 sequences (Fig S7). The CYP107 family includes *txtE* genes encoding nitrating enzymes that use NO as a substrate for the production of secondary metabolites and have no known role in respiratory denitrification or detoxification (40, 41). Thus, P450nor and TxtE are related (40, 41), yet TxtE is involved in SM and is the only other P450 observed to directly utilize NO as a substrate.

**Figure 2.**
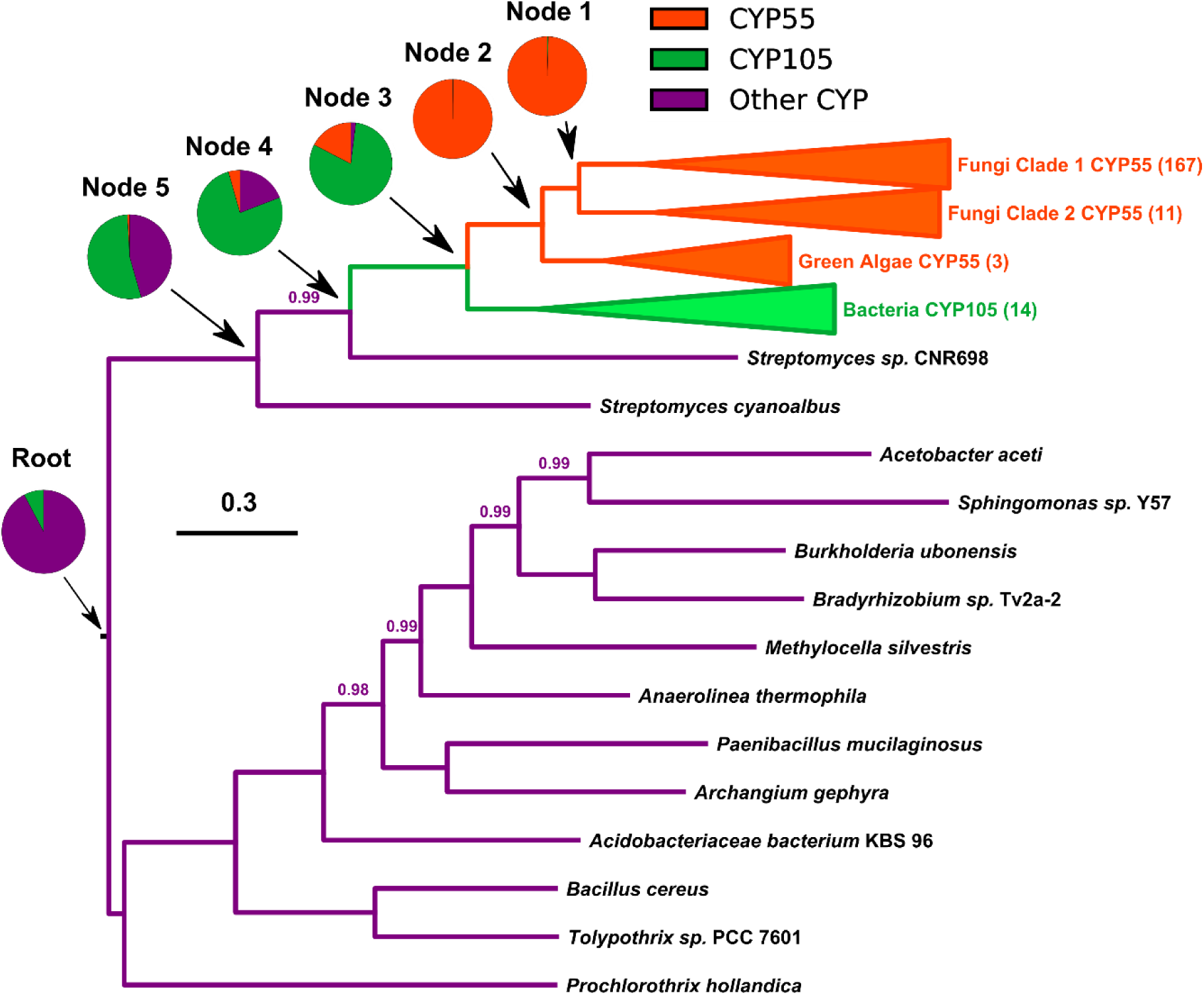
Midpoint-rooted Bayesian phylogeny of select families of cytochrome P450 amino acid sequences from fungi, algae, and bacteria. Ancestral state reconstruction was performed using CYP55 (orange), CYP105 (green), and other CYPs (purple) to uncover the shared ancestry of algal and fungal N2Ü-producing cytochrome P450s with their most recent common bacterial ancestor. The scale bar indicates substitutions per site and posterior probability values < 1 are displayed above branches of the Bayesian MCMC analysis. Numbers in parentheses next to collapsed clades indicate the number of sequences in the clade. Values in pie charts are average probabilities of each character state across one representative Bayesian MCMC analysis.

Sequences of the bacterial CYP105 family of P450s include diverse actinobacterial genera such as *Streptomyces* (n = 159), *Amycolatopsis* (n = 12), *Saccharothrix* (n = 5), *Streptacidiphilus* (n = 4), *Frankia* (n = 4), *Kutzneria* (n = 4), *Nocardia* (n = 3), and members from 17 additional actinobacterial genera (n = 39). The proteobacterial sequences were affiliated with members of the genera *Burkholderia* (n = 5), *Paracoccus* (n = 3), *Bradyrhizobium* (n=3), *Pseudomonas* (n = 1), and *Halomonas* (n = 1). Bacterial P450 gene and species tree comparisons of 60% identity clustered P450 amino acid sequences (n = 57) and cognate 16S rRNA genes (n = 55) supported HGT of one or more actinobacterial P450 genes to members of the *Alpha*-, *Beta*-, and *Gammaproteobacteria* (Fig. S6). Furthermore, ancestral character state reconstruction overwhelmingly supported *Actinobacteria* as the root state (root probability = 0.99 ± 0.06) of the bacterial CYP105 family P450 phylogeny. When forcing the root state of the P450 phylogeny to be *Proteobacteria* (simple model) and comparing to the complex model where the root is allowed to vary, the simple model with a proteobacterial root was not supported (average log Bayes Factor = 0.03 ± 0.18). Therefore, *p450nor* likely evolved from one or more CYP105 family P450 genes found in members of the *Actinobacteria.* This finding underscores *p450nor*’s distinct origin compared to the fungal denitrification traits *napA* and *nirK*, which have a distinct proteobacterial ancestry consistent with the majority of bacterial denitrifiers (Fig. S8).

### Widespread co-occurrence of *p450nor* and NO-detoxifying flavohemoglobins

Poor conversion of inorganic N-oxides to N_2_O by fungal isolates supports the hypothesis that P450nor is involved in NO detoxification (7, 42). However, fungi also possess NO-detoxifying flavohemoglobins responsible for detoxification of NO to NO_3_^−^ under oxic conditions or NO to N_2_O under anoxic conditions (42–44). Flavohemoglobins were detected in 450 (63 %) fungal genomes investigated and were widespread within ascomycete and basidiomycete fungi. Within *p450nor*-containing genomes, 125 (75 %) also possessed a flavohemoglobin gene (Fig. 1, Table S1). The number of genomes in fungal families containing *p450nor* and NO-detoxifying flavohemoglobin genes were significantly correlated (Spearman r = 0.87, *p* = 1.6e^−10^), and suggests *p450nor*’s primary function is not NO detoxification.

### Evidence of a role for *p450nor* in secondary metabolism

*p450nor* is actinobacterial in origin, yet *Actinobacteria* are not considered canonical denitrifiers and evidence for their role in denitrification was lacking when *p450nor* was initially identified (29, 45). Subsequent investigations did not posit a role for *p450nor* in SM despite the affiliation of *p450nor* and CYP105 P450s with documented roles in SM (34, 46). To assess genomic evidence linking *p450nor* to SM, we queried genes encoded within genomic regions approximately 50 kb on either side of *p450nor* for functions related to SM. The biosynthetic gene cluster (BGC) prediction tool antiSMASH detected putative BGCs containing *p450nor* in 19 (11 %) of the 167 *p450nor*-containing genomes analyzed (Dataset S2). The number of open reading frames in a predicted SM cluster ranged from 34 to 97, spanning 21,086 to 55,473 nucleotides in length. Inspection of protein-coding genes surrounding *p450nor* using curated antiSMASH profile Hidden Markov Models (pHHMs) resulted in the identification of hallmark SM features (e.g., polyketide synthases (PKS), non-ribosomal peptide synthases (NRPS), terpene cyclases, dimethylallyl tryptophan synthases) in an additional 40 (24 %) of the 167 *p450nor*-containing genomes analyzed (Dataset S2) (see Materials and Methods for details). The distribution of automatic and manually curated protein-coding genes surrounding a subset of 32 *p450nor*-containing fungi suggests that *p450nor*-containing BGCs are structurally and functionally diverse (Fig. 3). An additional BGC prediction tool, CASSIS, which detects BGCs based on shared transcription factor binding sites upstream and downstream of a user specified anchor gene (47), predicted as many as 105 (63 %) *p450nor*-containing gene regions to be BGCs (Dataset S3). Furthermore, CASSIS analysis corroborated 74 % of the 19 BGCs predicted by antiSMASH (Dataset S3). A detailed accounting of antiSMASH and CASSIS predictions, gene annotations, and gene organization surrounding *p450nor* in all 167 *p450nor*-containing genomes is available in the SI (Fig. S9, Dataset S2).

**Figure 3.**
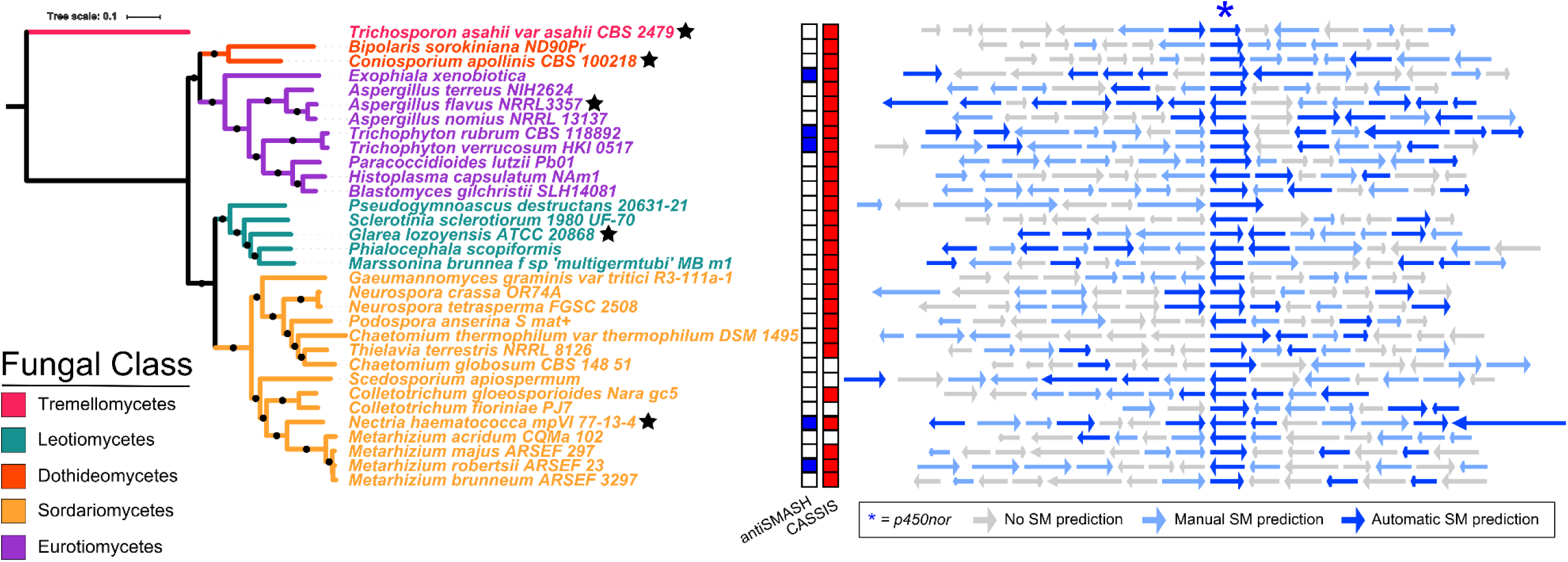
SM gene cluster predictions for a subset of 32 (167 total)*p450nor*-containing fungi. The boxes to the right of the rooted Maximum-Likelihood phylogeny indicate whether the *p450nor*-containing genomic region was predicted by antiSMASH (blue squares) or CASSIS (red squares) to be an SM gene cluster. White squares indicate no prediction. Colored arrows indicate protein-coding genes surrounding *p450nor* that were automatically predicted (dark blue arrow), manually predicted (light blue arrow) or not predicted (grey arrow) to be involved in SM (see Materials and Methods for details). The black stars next to species names are individuals chosen for in depth presentation of the genes surrounding *p450nor* (Fig. S9).

A diversity of secondary metabolite biosynthesis pathways were predicted to be encoded by *p450nor*-containing BGCs, including nonribosomal peptides (n = 7), polyketides (n = 5), terpenes (n = 2), hybrid terpene-polyketide-indoles (n = 2), indoles (n = 1), or currently unclassifiable compounds (n = 2). Phylogenetic reconstruction of C-type and ketosynthase domains encoded by NRPS and PKS genes surrounding *p450nor* enabled the prediction of potential secondary metabolites encoded by fungal *p450nor*-containing BGCs (Fig. 4). For example, domains from NRPS and PKS sequences encoded nearby *p450nor* are affiliated with reference NRPS and PKS sequences known to produce cyclic tetrapeptides (HC-toxins) (n=7), aflatoxins (n=5), fumonisins (n=4), calcium-dependent antibiotics (n=1), and statins (n=1), suggesting a large variety of secondary metabolites are encoded by gene regions containing *p450nor*.

**Figure 4.**
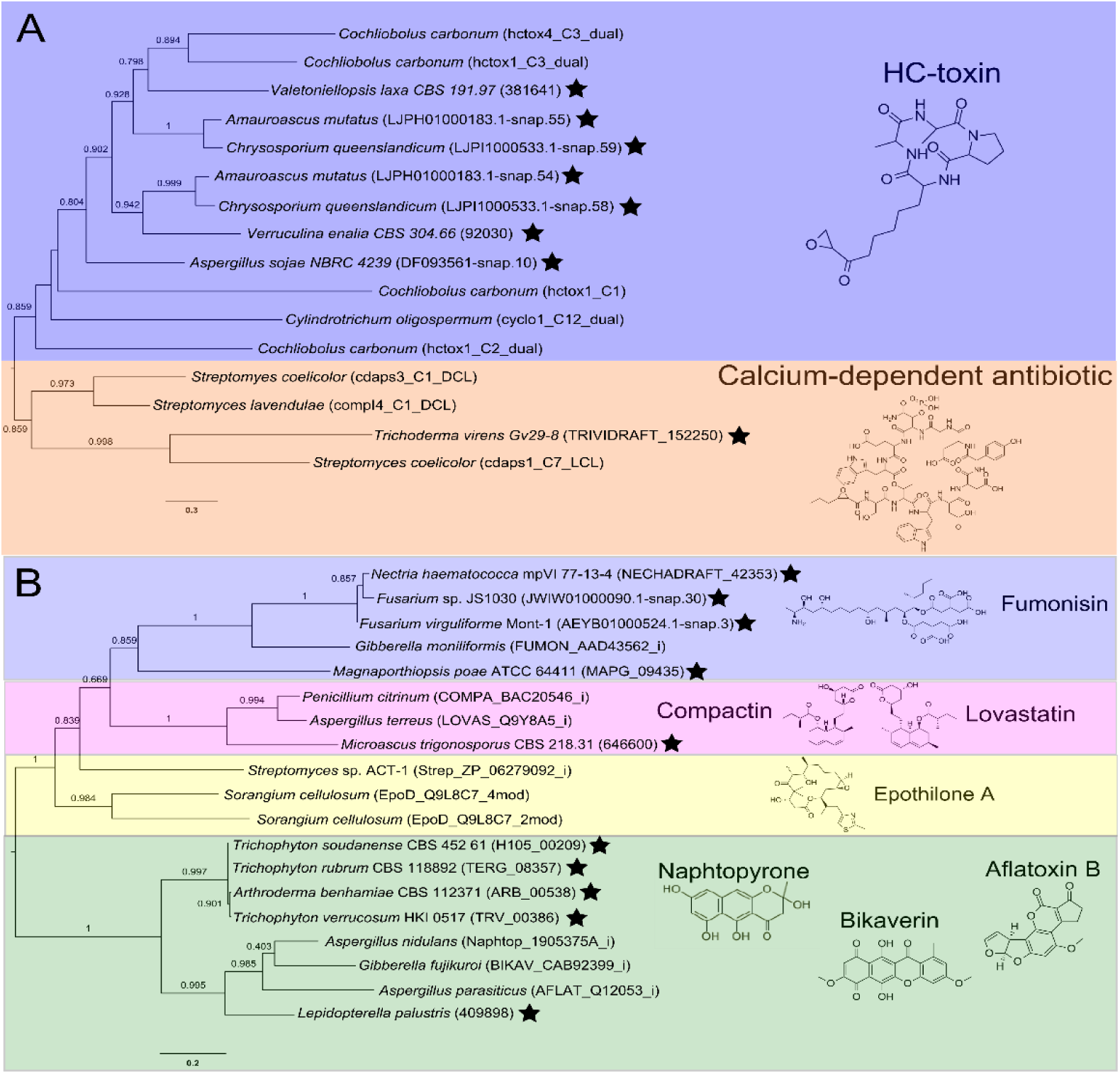
Maximum-Likelihood phylogenetic trees of non-ribosomal peptide synthase and polyketide synthase domains encoded within *p450nor*-containing genomic regions. Each phylogeny displays relationships of C-type condensation (C-type) (A) or ketosynthase (KS) domains (B) detected in non-ribosomal peptide and polyketide synthase amino acid sequences, respectively, encoded within *p450nor*-containing genomic regions. A black star next to taxa indicates C-type or KS domains identified in fungal genomes nearby *p450nor*. The NCBI or JGI accession numbers are shown in parentheses next to taxa with black stars. Taxa without black stars are reference amino acid sequences of C-type and KS domains curated by the NAPDOS database, and their NAPDOS accession numbers are indicated in parentheses. Proteins from species without accession numbers were predicted *ab initio* using SNAP (90) (see Materials and Methods for details). Chemical structures and names of secondary metabolites produced by NAPDOS reference sequences are indicated and highlighted distinct colors for clarity. Scale bars indicate amino acid substitutions per site. Values along branches indicate bootstrap support for the adjacent node.

The formation of N_2_O has previously been reported as highly variable among closely related fungi (37, 48), yet evidence suggesting a role for *p450nor* in this phenomenon is lacking. Of the 94 fungal genera harboring *p450nor*, 21 (22 %) contained species with and without a copy of *p450nor* (Table S5). For example, 15 out of 16 (94 %) *Pseudogymnoascus* genomes contained *p450nor*, whereas only 1 out of 7 (14 %) *Exophiala* genomes contained a *p450nor* gene. Nucleotide alignments of *p450nor*-containing genomic regions (81.3 ± 27.8 kb in length) against other fungal genomes revealed a disproportionately high nucleotide identity and alignment length between genomes with and without *p450nor* from the same genus (Fig. 5A-C). For example, genomic regions surrounding *p450nor* in *Exophiala xenobiotica* are highly similar to other *Exophiala* species without *p450nor* (Fig. 5D), and additional examples of large, high identity regions between closely related fungal genomes with and without *p450nor* are abundant (Fig. 5A-C, Dataset S4).

**Figure 5.**
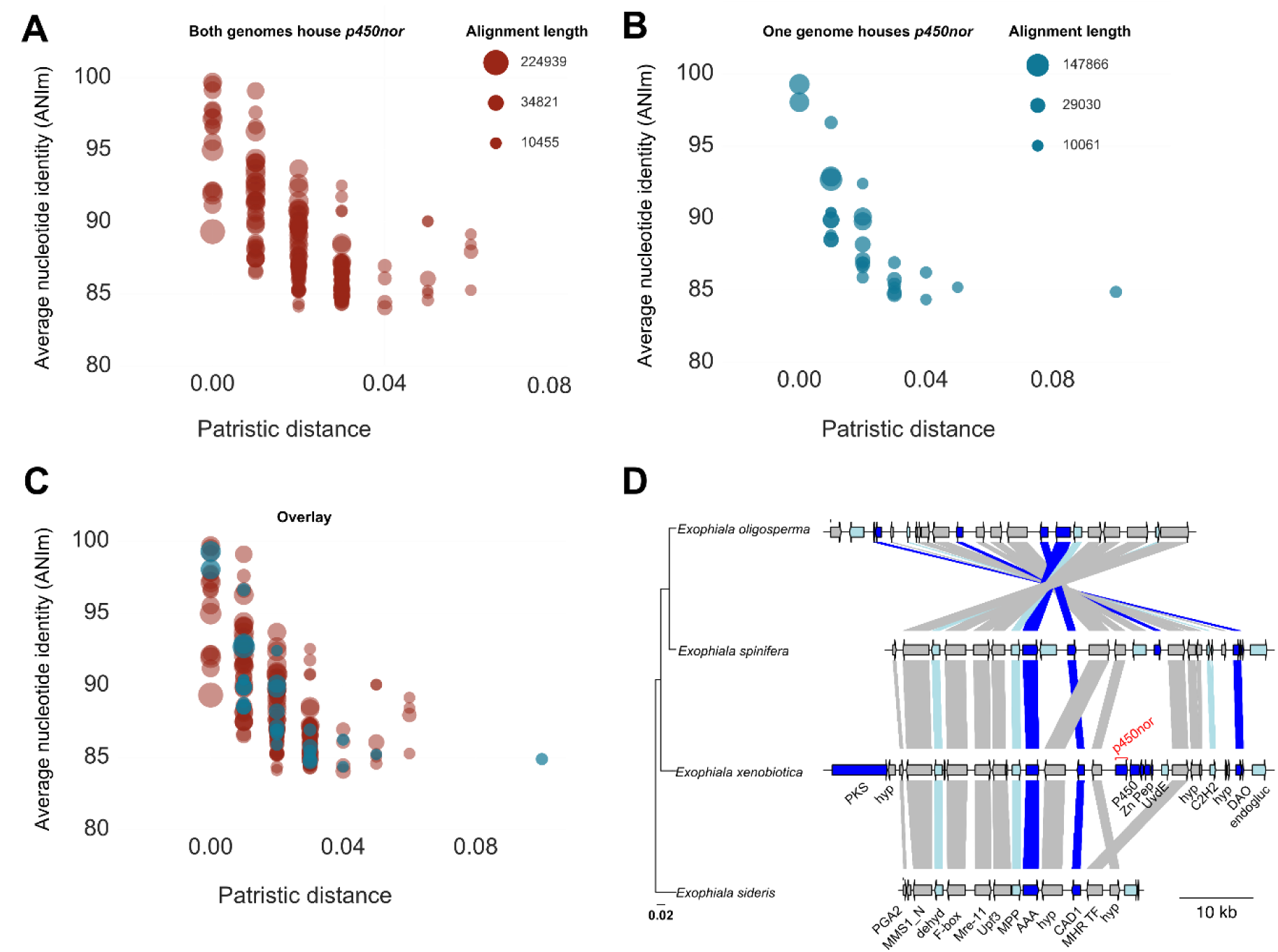
Within genera alignments of *p450nor*-containing genomic regions (N = 136) from species with and without *p450nor.* Alignment of two species with *p450nor*-containing genomic regions (red circles) (A) or between two species where only one member possesses *p450nor* (teal circles) (B). The overlay of the two plots in (C) indicates conservation of genomic architecture regardless of *p450nor* presence or absence. The size of the circles is proportional to the square root of the aligned length of the genomic regions. The gene synteny plot (D) highlights conservation of genomic architecture using a *p450nor*-containing genomic region from *Exophiala xenobiotia* aligned to three additional closely-related *Exophiala* species that do not possess *p450nor.* The arrows represent gene models within the gene region displayed. Both arrows and connecting lines above and below arrows are colored according to Figure 3 (see Materials and Methods for details). Lines connecting arrows between species indicate the genes are homologous. The scale bars (left to right) in (D) indicate substitutions per site and genome size in kilobases, respectively. The location of *p450nor* in *E. xenobiotica* is indicated in red font. The labels in black font describes the putative functions of proteins encoded by each gene. PKS - polyketide synthase, hyp – hypothetical protein, PGA2 – protein trafficking protein, MMS1_N – MMSl-like protein, dehyd – dehydrogenase, F-box – F-box domain containing protein, Mre-11 – double strand break repair protein Mre-11, Upf3 – nonsense mediated mRNA decay protein 3, MPP – metallophosphatase, AAA – ATPase, CAD1 – cinnamyl alcohol dehydrogenase, MHR TF – middle homology region transcription factor, P450 – cytochrome P450 reductase, Zn Pep – zinc peptidase superfamily protein, UvdE – UV-endonuclease, C2H2 – zinc finger C2H2 type, DAO – D-amino acid oxidase, endogluc – endoglucanase.

## Discussion

### An evaluation of hypotheses regarding the biological role of *p450nor*

The three leading hypotheses regarding the biological role of fungal *p450nor* are respiratory denitrification (7), NO detoxification (42), and now secondary metabolism. The respiratory denitrification hypothesis is dubious since evidence is lacking to classify fungi as respiratory denitrifiers (4, 17–19, 49). Furthermore, unaccounted for methodological biases inherent to partitioning techniques raises substantial concerns over the validity of fungal N_2_O production *in situ* (4, 25–27). The ineffectiveness of antibiotics to partition microbial respiration has been previously demonstrated (25, 26), yet antibiotics continue to be used to support the prevalence of fungal respiratory denitrification. In addition to antibiotic inhibition, site preference measurements of the intramolecular distribution of ^15^N within the linear N_2_O molecule (i.e., N_2_O isotopocules) of cultured microorganisms have been increasingly applied to partition microbial sources of N_2_O *in situ* (5, 50). Although promising, the limitations of N_2_O isotopocule measurements used in isolation are becoming apparent (27, 51, 52). Of primary concern is the significant overlap in, and difficulty discretizing, site preference measurements of distinct processes or diverse microbial assemblages (51, 53, 54). Therefore, the respiratory denitrification hypothesis is predicated on biased approaches used in isolation that are unable to correctly assess fungal contributions to denitrification.

Another alternative function suggested for P450nor is NO detoxification, which was initially postulated in experiments using the fungus *Fusarium oxysporum* strain 11n1 (55). This hypothesis was supported by low growth yields and a poor mass balance between the N-oxyanion inputs and N_2_O formed (18, 56, 57). Although plausible, the NO detoxification hypothesis is confounded by extensive co-occurrence between *p450nor* and genes encoding canonical NO-detoxifying flavohemoglobins, which also produce N_2_O (44, 58) (Fig. 1, Table S1). Considering the extensive overlap in *p450nor* and flavohemoglobin gene presence (Fig.1), the utility of site preference values derived from N_2_O formed by fungi in pure culture is questionable. Furthermore, P450nor and flavohemoglobins would likely compete for NO under anoxic conditions, and experiments teasing apart their contributions to N_2_O formation are necessary to support the postulated role of P450nor in NO detoxification. The reported Michaelis constant (K_m_) of NO binding to P450nor ranges from 0.1 to 0.6 mM (11, 12) and is orders of magnitude higher than the 0.1 to 0.25 μM K_m_ reported for flavohemoglobins (59), suggesting flavohemoglobin would outcompete P450nor for NO binding. Hence, the higher affinity of flavohemoglobins for NO and their greater distribution in fungi would suggest a limited role for P450nor in NO detoxification (Table S1). Though fungi certainly detoxify NO, insufficient evidence exists to attribute this activity to P450nor.

The SM hypothesis has traction considering that P450nor is derived from CYP105 P450s (Fig. 2), all of which share a functional role in SM (34, 46, 60). Thus, the adaptation of P450nor to a novel niche in NO reduction and denitrification is unlikely. A more parsimonious hypothesis is that P450nor has maintained a role in SM as observed for related actinobacterial enzymes.

When *p450nor* was originally described, members of the *Actinobacteria* (e.g., *Streptomyces*) were already well established secondary metabolite producers and their N_2_O production was attributed to detoxification (45, 61). The monophyly of P450nor with the SM enzyme TxtE, the only other NO-utilizing P450, provides additional *a priori* support for P450nor’s role in SM (Fig. S7). P450nor’s role in SM is further corroborated by SM prediction tools where a sizeable proportion (35 %) of gene regions surrounding *p450nor* contained genes predicted to encode hallmark SM functions, and as many as 105 (63 %) *p450nor*-containing genomic regions were automatically predicted to be involved in SM (Fig. 3). Moreover, the fact that antiSMASH flagged 11% of *p450nor*-containing genomic regions as putative BGCs suggests their organization and gene content is highly similar to other characterized BGCs. Although phylogenomic evidence supports a role for P450nor in the biosynthesis of secondary metabolites, direct physiological evidence should be a target for future research efforts. Emerging technologies enabling the expression of full length BGCs and metabolite identification should enable robust experimentation to test the SM hypothesis (62). Regardless, *p450nor*-containing genomic regions were predicted to be BGCs encoding diverse metabolites including terpenoids, nonribosomal peptides, polyketides, indoles, and other complex metabolites (Fig. 4) consistent with its evolutionary origins (Fig. 2).

### Predicting P450nor’s role in secondary metabolism

A variety of metabolites containing nitro functional groups have been detected in fungal genera known to harbor denitrifying representatives (63), yet mechanistic explanations for nitration reactions in fungi remain elusive. The addition of a nitro functional group to a metabolite represents a potential mechanism for enhancing its toxicity or functional specificity (64). The hypothesis of a role for P450nor in nitration, or possibly nitrosylation, of fungal metabolites is attractive given P450nor’s affiliation with the nitrating enzyme TxtE. The inclusion of *p450nor* within BGCs may be adaptive in fungal lineages in which this gene was acquired due to the augmenting effects that nitro or nitroso groups impart on their substrates (64). Support for this hypothesis stems from the widespread distribution of *p450nor* within secondary metabolite producing members of the Ascomycota (65, 66), and previous reports of HGT between members of *Actinobacteria* and fungi in enhancing fungal SM (67). Furthermore, the high nucleotide identity shared between *p450nor*-containing genomic regions from closely related fungal species suggests *p450nor* gain or loss may have important consequences for the secondary metabolites potentially produced by *p450nor*-containing BGCs (Fig. 5D). Considering that 22 *p450nor* containing fungal genera display variability in *p450nor* presence/absence (Table S5), investigations regarding the impact of *p450nor* presence/absence on the secondary metabolite pool, fungal fitness, competition, or infectivity within closely related fungi is readily testable.

Additional unknowns related to P450nor’s role in SM are the identification of putative substrates and sources of NO required to fuel the hypothesized nitration or nitrosylation reactions. To date, P450nor is solely reported to bind the electron donors NADH or NADPH and the electron acceptor NO (7). However, N_2_O formation by P450nor is oxygen dependent (8, 22), suggesting O_2_ may be an additional substrate as observed for TxtE (40). TxtE and NovI, both P450s affiliated with P450nor, bind to and transform L-tryptophan and L-tyrosine to produce the secondary metabolites thaxtomin A and novobiocin, respectively (40, 68). It is conceivable that P450nor might also bind O_2_ and aromatic amino acids, but direct experimental evidence is required to support this hypothesis. A potential source of NO in fungi could result from nitrite reductase activity of the copper containing nitrite reductase, NirK. The NO synthase (TxtD) from *Streptomyces turgidiscabies* produces NO to fuel TxtE nitration of L-tryptophan (40), but *txtD* homologs were not detected in the fungal genomes examined. Although evidence of NO synthases in fungi exist, knowledge regarding their distribution is limited (69, 70). Given the functional redundancy between NO synthases and NirK, it is conceivable that one of NirK’s functions in fungi is to generate NO for use by P450nor in SM.

### Causes and consequences of *p450nor* evolution in fungi

A limited understanding of *p450nor* evolution represented an impediment to our knowledge of fungal N_2_O formation. For example, closely related fungi vary in their ability to produce N_2_O (16, 37, 48, 56), and the evolutionary forces (e.g., HGT, gene gain/loss, and incomplete lineage sorting) contributing to this observation were unexplored. For *p450nor*, many HGT events were observed between distantly related fungal lineages using gene and species tree comparisons (Fig. S2). Although HGT events are challenging to precisely quantify given the level of uncertainty in deeply branching nodes of the functional gene trees reported here, a signal of potentially double digit HGT events were observed using gene tree-species tree reconciliation (Table S3). Moreover, genetic elements encoding *pogo* family transposases (N = 9), retrotransposons (N = 4), and reverse transcriptases (N = 1) were in some cases detected adjacent to *p450nor* and may act as vehicles for dissemination of *p450nor* within fungi and between fungal chromosomes (Dataset S2).

N_2_O production was previously coined a widespread trait in fungi (37), yet genomic analysis suggests fortuitous N_2_O formation by fungi is largely restricted to members of the Ascomycota. For example, of the167 *p450nor*-containing fungal genomes identified, 163 were affiliated with members of the Ascomycota and only four with members of the Basidiomycota. N_2_O production has been reported for fungal isolates assigned to the recently revised phylum Mucoromycota (4, 71), yet no evidence of genes underlying denitrification were detected in available genomes from members of this phylum (Fig. 1). Denitrification markers were also absent from ascomycete yeast genomes (i.e., *Candida*, *Yarrowia*), though a number of N_2_O-producing ascomycete yeasts have been reported (56). Even within the Basidiomycota, N_2_O formation is restricted to a few taxa within the Tremellomycetes and Agaricomycetes (4), and at least for members of the Tremellomycetes, was likely the result of HGT from one or more members of the Ascomycota (Fig S2). The finding that genomes from fungi (e.g., ascomycete yeasts) previously observed to produce N_2_O did not possess denitrification traits was unexpected and suggests that experimental artifacts or other mechanisms, such as the NO-detoxifying activity of flavohemoglobins, may also contribute to N_2_O formation in fungi. In addition to fungi, species of green algae have been reported to produce small quantities of N_2_O, the production of which could, at least in part, be attributed to the presence of *p450nor* within this lineage (72–74). Despite these findings, green algae lack a mass balance between the inorganic N added and the N_2_O formed (74) and display low rates and quantities of N_2_O production on par with fungi (72), suggesting that N_2_O formation is not a respiratory process in these organisms. Considering the lack of evidence of respiratory denitrification in green algae and genomic evidence linking *p450nor* to SM in fungi, the SM hypothesis is an attractive explanation for the presence of *p450nor* in green algae as well.

*p450nor* genes within fungi also have implications for fungal pathogenesis (4). At least for some bacteria (e.g., *Neisseria*, *Brucella*, *Mycobacterium*), the presence of denitrification genes has been demonstrated to enhance virulence or detoxification of N-oxides produced by the host (75). Although the impact of denitrification gene acquisition on fungal pathogenesis is not well established, there is growing evidence for P450nor involvement in fungal virulence (4, 7). For example, *p450nor* gene expression is linked to *Fusarium* wilt in banana and cotton plants, yet mechanistic explanations of P450nor’s function during plant infection are lacking (76, 77). Notably, more than half of all *p450nor*-containing fungal species are known plant pathogens (4), and the involvement of *p450nor* in SM is consistent with and would support the plant pathogenic life history strategies of many *p450nor*-containing fungi.

The diversity of denitrifying microorganisms and the modularity of the pathway has led to the view of denitrification as a community function (78–80). Therefore, limited co-occurrence and correlated evolution between *napA*, *nirK*, and *p450nor* might suggest mutualistic interactions occur between fungal or bacterial species performing denitrification. However, gene co-occurrences and evolutionary correlations should be interpreted with caution as additional factors (e.g., shared ecological niche, selection pressures) related to fungal life history strategies may explain their distribution equally well. For example, N_2_O-producing fungi are frequently detected in, and cultivated from, highly disturbed, N-amended agricultural soils (4, 9, 16) and detoxification or N-oxide utilization traits may merely co-occur more frequently due to selection imposed by episodic N addition. Fungi also contain genes homologous to bacterial denitrifiers, but their presence does not guarantee a role in respiratory denitrification. For example, the presence of genes homologous to the bacterial NO reductase (*norB*) is not sufficient evidence for respiratory denitrification potential in bacteria (17, 75). The same is true of the abundant *napA* gene homologs detected in fungal genomes, which would suggest a robust capacity of fungi to perform dissimilatory nitrate reduction. Yet this is not the case, and many fungi only produce N_2_O when NO_2_^−^ is present (4, 18, 56).

In summary, fungi often produce little or no gaseous N from reduction of N-oxyanions and do not grow proportionally to the quantity of N-oxyanions consumed; thus, fungi cannot be classified as respiratory denitrifiers (17). Given the limited accounting of methodological bias in the study of N_2_O production by fungi (25–27), alternative explanations for the biological function of *p450nor* in fungi are likely and raises concerns over the validity of these techniques in estimating fungal contributions to N_2_O emissions. Although the P450nor NO detoxification hypothesis is plausible, available data are insufficient at present to definitively support a role for P450nor in this process. Considering that many canonical denitrifying fungi are also plant disease causing secondary metabolite producers and agricultural pests, the affiliation of *p450nor* with non-denitrifying actinobacterial sequences involved in SM and their inclusion in BGCs strongly endorses a biological role for *p450nor* in SM.

## Materials and Methods

### Datasets

Draft and complete fungal, algal, and bacterial genomes were accessed from the National Center for Biotechnology Information and the Joint Genome Institute on March 16^th^, 2016 and downloaded from their respective database utilities. A list of fungal, algal, and bacterial genomes and their taxonomic and database affiliations can be found in the Supplemental Information (SI) (Dataset S5).

### Gene marker identification

To identify gene markers within fungal genomes suitable for phylogenetic analysis, a database of 1,438 amino acid sequences of fungal single copy orthologs from the BUSCO tool v1.1b (81) were provided as queries to the genblastG search tool v1.0.138 (82). The genblastG tool performs amino acid alignment of protein queries against a six frame translated nucleotide subject sequence (genome) to find significant alignments and uses heuristic analysis to piece the appropriate gene models back together from high-scoring segment pairs identified using BLAST (83). Of the BUSCO gene models queried, 238 were used for phylogenetic tree reconstruction and were annotated using PfamScan against the Pfam A database and blastp against the uniprot database with default settings (84–87) (Dataset S6). The genblastG tool was also used to detect gene sequences involved in denitrification (NapA, NarG, NirK/NirS, NorB, P450nor, NosZ) from curated bacterial proteins in the FunGene repository (88) or proteins involved in NO detoxification (flavohemoglobins) identified in the literature (44). Denitrification gene models used in downstream phylogenetic analysis were manually curated against full length fungal reference sequences to ensure that accurate gene models were predicted for each organism in which the gene was detected. After identification of these genes in fungal genomes, alignment of the fungal NapA, NirK, NarG, and P450nor amino acid sequences with blastp against the plant, archaea, bacteria, protozoa, and fungi RefSeq protein databases (89) was performed to identify similar sequences in each taxonomic group. Protein sequences demonstrating significant alignment (≥ 60 % query coverage and ≥ 35 % amino acid identity) to fungal proteins were used in subsequent phylogenetic reconstructions.

### Gene prediction for comparative genomic analyses

The *ab initio* gene predictor SNAP (90) was used to predict gene models in fungal genomes where no such information was available (e.g., some draft genomes). In this case, one or several closely related fungal genomes containing gene models were selected based on phylogenetic affiliation to train SNAP for gene prediction. Although this methodology is limiting when closely related genomes are unavailable, gene models from close relatives were available for *p450nor*-containing genomes lacking gene predictions.

### Alien index calculations

The alien index (AI) was calculated as previously described and modified for use with a single gene (44). Briefly, pairwise amino acid sequence alignments were performed using blastp for fungal NapA, NirK, and P450nor sequences. The in group was defined as the aligned sequence with the highest bitscore (excluding the query) belonging to the same taxonomic class as the query sequence. Accordingly, the out group was defined as the aligned sequence with the highest bitscore not belonging to the same taxonomic class as the query. The maximum bitscore was the bitscore derived from the alignment of the query to itself. Therefore, AI is calculated as follows:

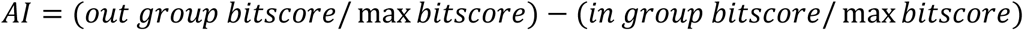

AI values range from 1 to −1. Values greater than zero are indicative of HGT or contamination of foreign DNA within the genome sequence being queried.

### Analysis of SM gene clusters in fungi

Genomic regions 50 kb up- and downstream of a *p450nor* gene in each genome were subjected to gene cluster prediction with the antiSMASH and CASSIS tools with default settings (47, 91). Additionally, genes encoded +/− 10 genes up and downstream of *p450nor* were evaluated using PfamScan searches with default settings against the pHHMs of curated SM genes identified by antiSMASH (Dataset S2) (91). Protein sequences with significant alignment to antiSMASH pHMMs were given an “automatic” SM function status and were colored blue. In order to supplement the automated SM annotation, additional functional annotation was performed by hmmscan searches with HMMER3 (92) against the eggNOG database (93). These functional annotations were manually flagged as related to SM if they possessed literature entries suggesting an involvement in SM or had functions related to methyl transfer, oxidation-reduction reactions, glycosyl transferases, fungal specific transcription factors, and other protein functions that may be important for SM outlined by antiSMASH (91). All manual SM annotations were colored light blue to indicate potential involvement in SM. All other annotations were colored grey when no evidence connecting the function to SM could be identified.

Additionally, ortholog clustering of protein-coding genes surrounding *p450nor* was performed using OrthoFinder (94) with default settings. Ortholog clustering was performed using only a representative *p450nor* loci in each fungal genome if multiple gene copies were present. A Shannon-like diversity index of fungal classes detected in each orthologous group was calculated as *H*’ = –Σ(*P_i_* ∗ ln(*P_i_*)), where *P_i_* is the fraction of fungal class *i* present in an orthologous group. Pairwise nucleotide alignments of *p450nor*-containing genomic regions were performed as previously described (95). Briefly, the *nucmer* utility of MUMmer *v3.0* (96) was used to align *p450nor*-containing genomic regions (∼100 kb) against whole genomes of fungi with and without *p450nor.* The average nucleotide identity (reported as ANIm) of the alignment was calculated from the resulting delta output file. The resulting data was plotted using Matplotlib (97) available for the python programming language (http://www.python.org).

### Phylogenetic analysis

Phylogenetic reconstruction of the fungal species tree was performed using concatenated amino acid sequences from 238 single copy orthologs found in ≥ 90 % of all genomes (Dataset S6). The genomes of *Puccinia arachidis* and *Microbotryum lychnidis*-*dioicae* strain p1A1 Lamole were excluded from further analysis due to an insufficient number of informative sites and inconsistent placement within the fungal tree. Alignment of amino acid sequences were performed individually on all 238 individual BUSCO gene models present within each organism using MAFFT v7.130b (98) with linsi alignment tuning parameters (‐‐maxiterate 1000 and ‐‐ localpair settings used). Individual alignments were concatenated using in-house python scripts, resulting in a 65,897 column alignment. Tree reconstruction was performed using FastTree2 (99) with refined tree reconstruction settings for slower, more exhaustive search of the tree space than default settings (-bionj -slow -gamma -spr 4 -lg -mlacc 2 and -slownni settings). For comparison to tree reconstruction using a concatenated alignment, individual trees from each BUSCO alignment were also constructed using FastTree2 with identical settings as above. The resultant alignments and trees were subjected to coalescent tree reconstruction using ASTRAL-II software (100). Overall, both phylogenies largely agreed except for branching patterns of some lineages (e.g., Zoopagomycota and Mucoromycota) and are available online in a figshare repository (see *Data Sharing* below)

The predicted amino acid and intronless nucleotide sequences of fungal *napA*, *nirK*, and *p450nor* gene models were aligned using the MAFFT settings described above and manually refined in JalView and SeaView software (101, 102). Maximum-likelihood (ML) and Bayesian phylogenetic tree reconstruction was performed on both nucleotide and amino acid alignments using RAxML and MrBayes, respectively (103, 104). Selection of the optimal evolutionary model for ML tree reconstruction was performed using prottest (105) (amino acid alignment) and jmodeltest (106) (nucleotide alignment) software prior to ML tree reconstruction. Please refer to SI for additional details about evolutionary models used in phylogenetic analysis.

Phylogenetic analysis with RAxML was performed by sampling 20 starting trees and performing 1,000 replicate bootstrap analyses. The tree with the maximal negative log likelihood score was compared to 1,000 replicates in RAxML to generate the final tree. Bayesian tree construction was performed using 3 independent runs with 6 chains for 5,000,000 generations. Output from MrBayes was evaluated with the sump and sumt commands within the software to ensure Markov Chain Monte Carlo chain mixing and convergence (potential scale reduction factor of 1.0) and standard deviation of split frequencies ∼ 0.01 or lower. MrBayes output was further visualized in the program Tracer (http://tree.bio.ed.ac.uk/software/tracer/) to ensure convergence was reached.

BayesTraits software was used to perform phylogenetically informed correlations between binary traits (i.e., the presence or absence of two denitrification markers) and ancestral state reconstruction (107). Please refer to SI for additional details on BayesTraits analyses.

Approximately unbiased (AU) tests were performed in the program CONSEL (108) using default settings. The negative log likelihood values from the observed nucleotide phylogenies input into CONSEL were −140,261, −37,782, −111,531 for *napA*, *nirK* and *p450nor*, respectively. The observed negative log likelihood scores for amino acid phylogenies of NapA, NirK, and P450nor were −65,158, −14,197, and −44,171, respectively. Species-tree gene-tree reconciliation was performed using NOTUNG software v2.9 (38, 109). Please see SI for further details on NOTUNG parameters.

### Statistical analysis

All statistical analyses were carried out in R programming language (110) and significance of statistical tests were assessed using a *p* value cutoff ≤ 0.05.

### Data sharing

All gene models, alignments, and trees discussed in the manuscript are made available in a figshare repository prepared by S.A.H. (https://doi.org/10.6084/m9.figshare.c.3845692).

## Acknowledgements

We thank Gerald Bills, Gregory Bonito, Pedro Crous, Kathryn Bushley, Colleen Hansel, Patrik Inderbitzin, Gabor Kovacs, Bjorn Lindahl, Jon Magnuson, Francis Martin, Kerry O’Donnell, Nancy Nichols, Minou Nowrousian, and Joseph Spatafora for providing access to unpublished genome data produced by the U.S. Department of Energy Joint Genome Institute. The authors thank A. Frank for assistance with manuscript revisions. S.A.H. would like to acknowledge a Department of Energy Office of Science Graduate Student Research fellowship for support during preparation of the manuscript. Participation of C.W.S and F.E.L. was partially sponsored by the Laboratory Directed Research and Development Program of Oak Ridge National Laboratory, managed by UT-Battelle, LLC, for the U. S. Department of Energy. This research was partially supported by the US Department of Energy, Office of Biological and Environmental Research, Genomic Science Program, Award DE-SC0006662.

## List of Supplemental Materials

**Table S1**. Counts of denitrification traits and their co-occurrences in fungal genomes.

**Table S2**. Results from approximately unbiased tests for the monophyly of fungal classes within *napA*, *nirK*, and *p450nor* gene trees. Where indicated, the monophyly of two lineages was also assessed. Bold font data indicate that the AU test rejected the monophyly of the taxa. Test significance was evaluated at *p* ≤ 0.05.

**Table S3**. Results from species-tree gene-tree reconciliation using NOTUNG software for *napA*, *nirK*, and *p450nor* genes in fungi. Values are averages of solutions with standard deviations reported in parentheses.

**Table S4**. Predicted horizontal gene transfers of fungal *p450nor*, *napA*, and *nirK* genes based on alien index algorithm.

**Table S5**. List of genera containing species with and without *p450nor.*

**Figure S1**. Gene abundances of *narG*, *napA*, *nirK*, *p450nor*, and flavohemoglobins (colored bars) mapped on to fungal families (cladogram, left). Relationships among fungal families in the cladogram were derived from the NCBI taxonomy using the online tool phyloT (http://phylot.biobyte.de/index.html).

**Figure S2**. Maximum-Likelihood phylogenies connecting fungal species with their respective NO reductase (*p450nor*) gene sequence(s). On the left, an amino acid phylogeny of 238 concatenated single copy orthologues from fungal species in which one or more *p450nor* gene(s) were detected. The *p450nor* nucleotide phylogeny (right) demonstrates many instances of incongruence with the fungal species phylogeny. Black dots in each phylogeny represent bootstrap percentages greater than or equal to 90%. Scale bars represent amino acid (left tree) and nucleotide (right tree) substitutions per site. A high-resolution file of the tree is available at https://doi.org/10.6084/m9.figshare.c.3845692.

**Figure S3**. Cophylogenetic plot of *napA*-containing fungal species (left, N = 75) and the *napA* nucleotide tree (right, N = 78). Both are midpoint rooted Maximum-Likelihood trees where black dots represent bootstrap percentages ≥90 %. Scale bars indicate substitutions per site for the concatenated amino acid species phylogeny and nucleotide phylogeny, respectively. A high-resolution file of the tree is available at https://doi.org/10.6084/m9.figshare.c.3845692.

**Figure S4**. Cophylogenetic plot of *nirK*-containing fungal species (left, N = 82) and the *nirK* nucleotide tree (right, N = 83). Both are midpoint rooted Maximum-Likelihood trees where black dots represent bootstrap percentages ≥90 %. Scale bars indicate substitutions per site for the concatenated amino acid species phylogeny and nucleotide phylogeny, respectively. A high-resolution file of the tree is available at https://doi.org/10.6084/m9.figshare.c.3845692.

**Figure S5**. Plot of alien index values observed for *p450nor* genes (N = 178). Points above the hashed line at the origin are indicative of HGT. Names of fungal species with alien index values above zero are ordered as their points appear on the graph. Thick horizontal lines represent the median alien index value. See Materials and Methods in the main text for details on alien index calculations.

**Figure S6**. Bayesian tree reconstruction of actinobacterial and proteobacterial 16S rRNA genes (left, N = 55) and cytochrome P450 family 105 amino acid sequences (right, N = 57). Both phylogenies represent 50% majority-rule consensus trees. The tree on the left is rooted with proteobacterial sequences as outgroup to the *Actinobacteria.* The tree on the right is midpoint rooted. Nodes with posterior probabilites ≥ 0.95 are indicated by black circles on an adjacent branch.

**Figure S7**. Midpoint rooted Bayesian (left) and Maximum-Likelihood phylogenies (right) of cytochrome P450 sequences (N = 408) demonstrating the affiliation of P450nor with other sequences belonging to members of the bacterial phyla Actinobacteria and Proteobacteria. Cyanobacterial cytochrome P450 sequences were included as outgroups. Black squares on branches (left tree) indicate ≥0.95 posterior probability or ≥90 % bootstrap replication (right tree). The colored legend indicates the cytochrome P450 family specified by shared amino acid identity of ≥40 % (D.R. Nelson, Hum Genomics 4:59-65, 2009).

**Figure S8.** Bayesian and Maximum-likelihood phylogenies of NapA, NirK, and P450nor amino acid sequence homologs extracted from the RefSeq protein database. A high-resolution file of these trees are available at https://doi.org/10.6084/m9.figshare.c.3845692.

**Figure S9**. Genome regions chosen for in depth presentation of protein coding genes surrounding *p450nor* in predicted BGC regions. Labels above genes are functional annotations from alignments to the eggNOG database. NCBI gene loci accessions are labeled below each gene.

